# Nomenclature Errors in Public 16S rRNA Gene Reference Databases

**DOI:** 10.1101/441576

**Authors:** Kyle Lesack, Inanc Birol

**Affiliations:** Department of Ecosystem and Public Health, Faculty of Veterinary Medicine, University of Calgary; Professor, Medical Genetics, University of British Columbia

## Abstract

**Background:** Targeted gene surveys of the 16S rRNA gene have become a standard method for profiling the membership and biodiversity of microbial communities. These studies rely upon specialized databases that provide reference sequences and their corresponding taxonomic classifications, but few independent evaluations of the nomenclature used in the taxonomic classifications have been performed.

**Results:** Nomenclature data collected from the List of Prokaryotic names with Standing in Nomenclature, Prokaryotic Nomenclature Up-to-Date, and CyanoDB databases were used to validate the nomenclature contained in the taxonomic classifications in the Greengenes, RDP, and SILVA 16S rRNA gene reference databases. Between 82% and 97% of the genus annotations assigned to 16S rRNA gene reference sequences were deemed valid in the reference databases. Between 18% and 97% of the species annotations in Greengenes and SILVA were deemed valid. Misannotations included the use of metadata in place of taxonomic classifications, non-adherence to the binomial nomenclature, and sequences classified as eukaryote organelles or taxa.

**Conclusions:** The misannotations identified in public 16S rRNA gene databases call into question the reliability of research made using these resources. As targeted gene surveys depend on high quality marker gene databases, imed nomenclature accuracy will be necessary.

## Background

Targeted gene surveys are a popular approach for inferring the population structure of complex microbial communities. Phylogenetically informative loci, such as the 16S rRNA gene, are first amplified and sequenced. Community members may then be identified by mapping the sequencing reads from a sample to a reference database containing both the marker gene reference sequences and taxonomic annotations. Despite technical advances, errors introduced from molecular and computational methods remain a problem. Errors introduced during the PCR and sequencing stages are well described, and include amplification bias^1,2^, PCR chimeras^3,4^, and incorrect base calling^3,5,6^. Errors may also result from computational methods, such as operational taxonomic unit clustering^7,8^, copy number correction^9^, and taxonomic classification ^10^. Together, these errors can have a dramatic impact on the accuracy of taxonomic classifications and biodiversity estimates^7^.

Access to high quality reference data is crucial for marker gene surveys, as reference databases provide two key functions: they allow microbial communities to be profiled by mapping unknown sequencing reads to known organisms, and they provide reference sequences for multiple sequence alignments used in biodiversity calculations. Nucleotide sequences available in the public International Nucleotide Sequence Database Collaboration (INSDC) databases are usually annotated by the submitting author at the time of deposition with little quality control measures to ensure annotation accuracy, allowing for the propagation of error in subsequent analyses. Therefore, several specialized 16S rRNA gene databases have been curated with the aim of providing high quality reference data for microbial gene surveys. Three main databases that include broad coverage of 16s rRNA gene sequences are available for download: Greengenes^11^, the Ribosomal Database Project^12^, and SILVA^13^.

Despite the importance of high quality reference data for marker gene surveys, few quality assessments of the public 16S rRNA gene reference databases have been performed. One study, published in 2005, suggested that at least 5% of the sequences deposited in a previous release of the RDP database (release 9, update 22) contained sequence anomalies, such as PCR chimeras or base calling errors^14^. More recently, two studies have examined the quality of taxonomic annotations in Greengenes, RDP, and SILVA. One study estimated the levels of taxonomic misannotation in Greengenes, RDP, and SILVA at 0.2%, 1.27%, and 1.54% respectively^15^. Conversely, the other estimates suggested that they were as high as 10% for RDP, and 17% for both Greengenes and SILVA^16^.

The use of invalid nomenclature is an underappreciated type of misannotation in biological databases. While nomenclature is closely allied to taxonomy, the rules governing the naming of organisms are separate from taxonomic interpretations. Therefore, estimates of taxonomic misannotations should not be considered as indicative of the quality of nomenclature annotations. Database annotations may be taxonomically consistent, but at the same invalid according to the governing code of nomenclature. To be validly published, prokaryote names must meet the requirements described in the International Code of Nomenclature of Prokaryotes (ICNP; formerly the International Code of Nomenclature of Bacteria)^17^. The Approved List of Bacterial Names was published in 1980, designating which of the existing prokaryote names described in the literature would be considered valid and retained for the future. Following the publication of the Approved List of Bacterial Names, publication in the International Journal of Systematic and Evolutionary Microbiology has been a requirement for valid publication.

The regulation of cyanobacterial nomenclature remains unsolved. Historically, cyanobacteria were identified as algae, and governed under the International Code of Nomenclature for Algae, Fungi, and Plants (ICN). However, phylogenetic analyses have revealed that cyanobacteria should be classified as bacteria^18^. Because the rules governing nomenclature differ between the ICNP and ICN, there is a need to standardize the nomenclature of cyanobacteria. Two dissimilar proposals for the regulation of cyanobacterial nomenclature have been submitted to the Special Committee on Harmonization of Nomenclature of Cyanophyta/Cyanobacteria. The first proposal suggests excluding cyanobacteria from the ICNP^19^, while the later proposal argues for applying the ICNP rules to all cyanobacteria^20^. At present, neither proposal has been adopted^21^.

A thorough validation of the nomenclature contained in 16S rRNA gene databases is challenging due to limitations of the available resources on prokaryote nomenclature. Taxonomic classifications assigned to sequences uploaded to the INSDC databases are a primary source of nomenclature used in subsequent analyses, however, there is no quality assurance used to ensure that a given name is validly published. Moreover, the NCBI taxonomy group have stated that their taxonomy should not be considered as an authoritative resource (https://www.ncbi.nlm.nih.gov/Taxonomy/taxonomyhome.html/index.cgi?chapter=howcite).

Comprehensive lists of valid and invalid prokaryote names are available from the List of Prokaryotic Names with Standing in Nomenclature (LPSN)^22^ and Prokaryotic Nomenclature Up-to-Date ^23^ databases. For cyanobacteria, valid and invalid names are available from CyanoDB^24^. Collecting data from these resources presents a challenge, as the data are provided in formats suitable for the validation of individual prokaryote names, but infeasible for the validation of entire databases containing millions of records.

To address these challenges, we developed custom Python scripts to automate the collection of nomenclature data from LPSN, Up-to-Date, and CyanoDB. Using data collected from these resources, the nomenclature annotations contained in Greengenes, RDP, and SILVA were validated. For each database, a considerable proportion of the nomenclature annotations were either invalid or of unknown validity. Sequences classified as mitochondria or chloroplasts were found in all three databases, and sequences classified using eukaryote names were identified in RDP and SILVA. Names failing the binomial nomenclature were identified in SILVA.

## Results

### Prokaryote Nomenclature Validation Dataset

Valid and invalid prokaryote genus and species names were collected from CyanoDB, LPSN, and Prokaryotic Nomenclature Up-To-Date (Table 1). A validation set was created from these data to evaluate the reliability of nomenclature annotations in the 16S rRNA gene reference databases.

**Table 1.**
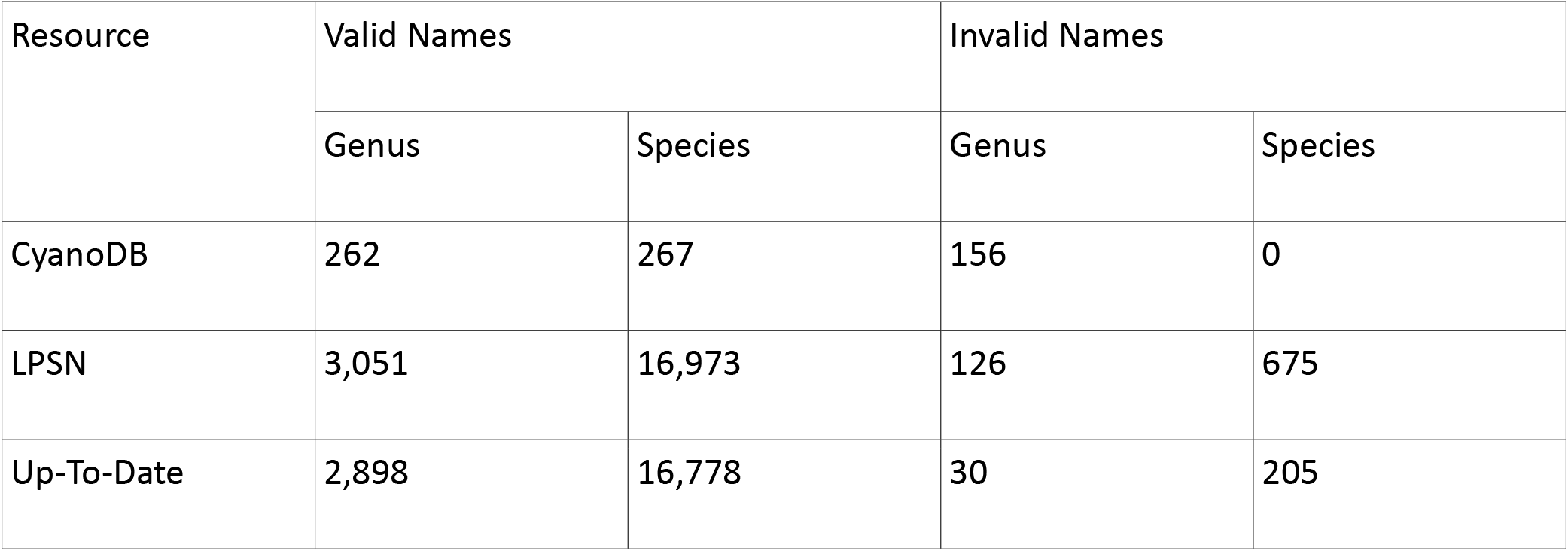
Valid and invalid genus and species names collected from prokaryote nomenclature reference databases.

| Resource | Valid Names |  | Invalid Names |  |
| --- | --- | --- | --- | --- |
|  | Genus | Species | Genus | Species |
| CyanoDB | 262 | 267 | 156 | 0 |
| LPSN | 3,051 | 16,973 | 126 | 675 |
| Up-To-Date | 2,898 | 16,778 | 30 | 205 |

Names were considered valid if they were deemed valid in at least one reference database and were not deemed invalid by any other. Similarly, names were considered invalid if they were deemed invalid in at least one reference database and were not deemed valid by any other. Disagreements on the validity of names between databases were categorized as disputed. The final validation set contained 3,282 valid, 246 invalid, and 63 disputed names at the genus level. 17,381 valid, 717 invalid, and 163 disputed names were collected at the species level.

**Table 2.** Final Validation Set.

| **Rank** | **Category** | **Count** |
| --- | --- | --- |
| Genera | Valid | 3,282 |
|  | Invalid | 246 |
|  | Disputed | 63 |
| Species | Valid | 17,381 |
|  | Invalid | 717 |
|  | Disputed | 163 |

### Validation of Nomenclature Contained in 16S rRNA Gene Reference Databases

Three specialized 16S rRNA gene databases (Table 3) were evaluated in this study: Greengenes (v. 13_5), RDP (release 11, update 5), and SILVA (SSU Ref, v. 132). The taxonomy files for each database were downloaded, and the genus and species (Greengenes and SILVA only) names assigned to reference sequences were evaluated using the validation set.

**Table 3.** Public 16S rRNA Gene Reference Databases.

| Name | Version | 16S rRNA Gene Sequences | Sequences Classified to the Genus Level | Sequences Classified to the Species Level |
| --- | --- | --- | --- | --- |
| Greengenes | 13_5; May 2015 | 1,262,986 | 878,985 | 285,288 |
| RDP | Release 11, Update 5; September 2016 | 3,356,808 | 2,394,396 | 0 |
| SILVA (SSURef)<sup>1</sup> | 132; December 2017 | 1,928,733 | 1,802,698 | 1,722,495 |

1 – Multiple SILVA databases are available. The present analysis is based on the SSUREF database

A manual inspection of the annotations assigned to reference 16S rRNA gene sequences revealed the presence of metadata in place of many taxonomic classifications. Therefore, lists of obvious metadata were collected and used to filter out database records that were annotated with metadata in place of genus or species names (Additional file 1: Table S1). All three databases contained annotations labelled as *Candidatus*, a category used to describe putative taxa which do not meet the requirements for valid publication under the Bacteriological code. RDP contained many genera categorized with the suffix ″incertae sedis″ (e.g., Subdivision3_genera_incertae_sedis), which describes taxa of uncertain placement. Although taxa classified as *Candidatus* are not considered to be validly published, its usage is considered acceptable for describing uncultured prokaryote species whose taxonomic placement has been determined^25^. Therefore, sequences classified using the *Candidatus* category were quantified separately from those deemed invalid. Greengenes and SILVA contained sequences annotated with higher ranks (e.g., *uncultured Bacteroides sp., Bosea genosp*.). Although these annotations may contain valid names, they were categorized as metadata, as they did not provide classifications at the specified rank.

To reduce the number of uncategorized annotations, *Candidatus, Incertae sedis*, and metadata were included as separate categories in the validation analysis (Table 4). Annotations not belonging the above categories were evaluated using the validation set. Nomenclature annotations left uncategorized included names not described as valid or invalid by any of the nomenclature databases and names of disputed validity between the nomenclature databases.

**Table 4.** Validation of genus names in 16S rRNA Gene Databases.

| Database | Total genera | Valid Names | Invalid Names | Candidatus / Incertae | Annotated with Metadata | Uncategorized<sup>1</sup> | Uncategorized (Disputed Validity)<sup>2</sup> | Total Uncategorized |
| --- | --- | --- | --- | --- | --- | --- | --- | --- |
| Greengenes | 879,081 | 849,416<br>(96.63%) | 1,611<br>(0.18%) | 6,977 (0.79%) | 94 (0.01%) | 18,991 (2.16%) | 1,992 (0.23%) | 20,983 (2.39%) |
| RDP | 2,394,396 | 2,111,586<br>(88.19%) | 899<br>(0.04%) | 37,377 (1.56%)<br>/<br>48,280 (2.02%) | 0 | 181,331 (7.57%) | 14,923 (0.62%) | 196,254 (8.20%) |
| SILVA | 1802698 | 1,474,317<br>(81.78%) | 3,574<br>(0.20%) | 30,880 (1.71%) | 168,770<br>(9.36%) | 117,155 (6.50%) | 8,002 (0.44%) | 125,157 (6.94%) |

1 – These annotations were not found in the validity set.
2 – These annotations were left uncategorized because their validity was disputed between the nomenclature databases.

The percentage of valid genus names ranged from 96.63% (Greengenes) to 88.19% (RDP) and 81.78% (SILVA). Few annotations matched to known invalid genera, as the percentage of invalid names ranged from 0.18% (Greengenes) to 0.04% (RDP) and to 0.20% (SILVA). These are likely conservative estimates, as many names were left uncategorized (Greengenes = 2.39%, RDP = 8.20%, SILVA = 6.94%). Names with disputed validity between the nomenclature databases only contributed to a small number of uncategorized annotations (Greengenes = 0.23%, RDP = 0.62%, SILVA = 0.44%). Considerable metadata was present in SILVA, as 9.36% of the records were annotated with metadata at the genus level. Only 0.01% of the genus annotations in Greengenes contained metadata. No metadata was identified in the RDP database genus annotations. Annotations labelled as *Candidatus* (Greengenes = 0.79%, RDP = 1.56%, SILVA = 1.71%) and *Incertae sedis* (RDP = 2.02%) accounted for the remaining records. The most common invalid and uncategorized genus annotations for each database are shown in the supplemental material (Additional file 1: Table S2).

97.26% and 18.14% of species names were categorized as valid in Greengenes and SILVA respectively (Table 5). Only 0.10% of the species names in Greengenes, and 0.08% of the species names in SILVA were classified as invalid. Again, these are likely conservative estimates, as the total uncategorized species names accounted for 2.02% of the species annotations in Greengenes and 0.76% of the species annotations in SILVA. Names with disputed validity between the nomenclature databases only contributed to a small number of uncategorized annotations (Greengenes = 0.03%, SILVA = 0.11%). SILVA contained a higher proportion of *Candidatus* species (1.85%) compared to Greengenes (0.45%), and considerably more species names annotated with metadata (SILVA = 79.19%, Greengenes = 0.18%). The most common invalid and uncategorized species annotations for each database are shown in the supplemental material (Additional file 1: Table S3).

**Table 5.** Validation of species names in 16S rRNA Gene Databases.

| Database | Total Species | Valid Names | Invalid Names | Candidatus | Annotated with Metadata | Uncategorized | Uncategorized (Disputed Validity)<sup>1</sup> | Total Uncategorized |
| --- | --- | --- | --- | --- | --- | --- | --- | --- |
| Greengenes | 285,289 | 277,462<br>(97.26%) | 280 (0.10%) | 1,274 (0.45%) | 511 (0.18%) | 5,687 (1.99%) | 75 (0.03%) | 5,762 (2.02%) |
| SILVA | 1,722,495 | 312,404<br>(18.14%) | 1405 (0.08%) | 31,783 (1.85%) | 1,363,770<br>(79.19%) | 11,160 (0.65%) | 1,973 (0.11%) | 13,133 (0.76%) |

1 – These annotations were not found in the validity set
2 – These annotations were left uncategorized because their validity was disputed between the nomenclature databases

### Eukaryote Annotations in 16S rRNA Gene Databases

Greengenes, RDP, and SILVA all contained eukaryote organelles as taxonomic groups (Table 6). In Greengenes 496 taxa were classified as belonging to chloroplasts, while 5,924 were classified as mitochondria. RDP contained 71,940 taxa classified as chloroplasts. 19,102 taxa in SILVA were classified as chloroplasts, and 2,265 as mitochondria.

**Table 6.** Sequences classified as mitochondria or chloroplasts.

| Database | Mitochondria | Chloroplasts | Total Organelles |
| --- | --- | --- | --- |
| Greengenes | 5,924 | 496 | 6,420 |
| RDP | 0 | 71,940 | 71,940 |
| SILVA | 2,265 | 19,102 | 21,367 |

Records misannotated using eukaryote names accounted for many of the annotations in RDP and SILVA that were not classified by the validation set (Table 7). Seven unique eukaryote taxa (Bacillariophyta, Bangiophyceae, Chlorarachniophyceae, Chlorophyta, Cryptomonadaceae, Euglenida, Streptophyta) accounted for 9.78% of the uncategorized genera in RDP. 835 unique eukaryote names accounted for 1.68% of the unclassified genera in SILVA (Additional file 2). 304 unique eukaryote names accounted for 6.42% of the uncategorized species annotations in SILVA (Additional file 3. No eukaryote names were found in the uncategorized Greengenes annotations.

**Table 7.** Sequences classified using eukaryote names.

| Database | Rank | Unclassified<br>Records | Unique<br>Eukaryote Taxa | Total Sequences Classified using<br>Eukaryote Taxon<sup>1</sup> |
| --- | --- | --- | --- | --- |
| GG | Genus | 18,991 | 0 | 0 |
|  | Species | 5,687 | 0 | 0 |
| RDP | Genus | 181,331 | 7 | 17,726 (9.78%) |
| SILVA | Genus | 117,155 | 835 | 1,965 (1.68%) |
|  | Species | 11,160 | 304 | 717 (6.42%) |

1 - Describes the percentage of unclassified sequences classified as eukaryote taxa

### Binomial Nomenclature in SILVA

SILVA contained 44,434 species annotations, where the genus epithet of the species names did not match the assigned genus. Half of the species annotations (22,250) failed the binomial nomenclature test due to the use of merged genus names to represent species belonging to non-monophyletic groups (e.g., *Escherichia coli* are placed in the *Escherichia-Shigella* genus^26^). SILVA contained 8,028 genus names with multiple space separated terms (e.g., *Microcystis viridis* was placed under the *Microcystis PCC-7914* genus), which accounted for 18% of the species names that failed the binomial nomenclature check. Complete mismatches (e.g., *Fluoribacter bozemanae* was placed under the *Legionella* genus) accounted for the remaining 14,156 (32%) species annotations that did not adhere to the binomial nomenclature. All Greengenes species annotations adhered to the binomial nomenclature.

## Discussion

Access to high quality 16S rRNA gene reference sequences and annotations are crucial for microbial ecology research. Although quality assurance methods were employed during the curation of the Greengenes, RDP, and SILVA databases, considerable misannotations were identified in the nomenclature contained in these databases. While the exact impact of these misannotations on the statistical calculations used in microbial gene survey projects is currently unknown, the results obtained here call into question the reliability of published work made using these databases.

Overall, the Greengenes database contained the fewest misannotations, as 97% of the genus and species names were deemed valid. Conversely, only 88% of the genera in RDP, and 82% of the genera in SILVA were deemed valid. The species names in SILVA were especially problematic, as only 18% of these annotations were deemed valid. Records annotated with metadata in place of taxonomic classifications were a major problem in SILVA, as metadata accounted for 9% and 79% of the annotations at the genus and species levels respectively. The invalid names collected for the validation set only included invalid names observed by the curators of the nomenclature databases and are not comprehensive lists of all invalid names in use. Therefore, the proportions of annotations that we classified as invalid are likely underestimates. We expect that most of the unclassified annotations are in fact invalid, rather than valid names missing from the nomenclature databases.

Annotations containing eukaryote taxonomic classifications and organelles were also identified in the 16S rRNA gene databases. Many prokaryote 16S rRNA primers have exhibited affinity for chloroplast and mitochondrial DNA^27^, and may explain how mitochondria and chloroplasts were included in public 16S rRNA gene databases. However, all these sequences were classified as bacteria, which is incorrect. Both RDP and SILVA included eukaryote taxonomic classifications. These misannotations may have occurred due to a sequence being classified using the host species name in place of the prokaryote species whose 16S rRNA gene was sequenced.

As microbial gene surveys continue to rely upon public 16S rRNA gene reference databases, a thorough review of the annotations contained in these databases will be required. The misannotations discussed above were identified using scripts that automate the collection of reference nomenclature and validation of the names contained in the Greengenes, RDP, and SILVA databases. These scripts are available to the public and may be useful for improved quality control for the curation of marker gene databases. Community awareness of these problems is also important. Users that contribute content INSDC databases need to be aware of how misannotations propagate errors and place further burdens on database curators with limited resources.

## Conclusions

This study assessed the validity of nomenclature annotations contained in the Greengenes, RDP, and SILVA databases. Considerable annotations were deemed as invalid or of unknown validity. All databases contained records annotated with metadata in place of taxonomic classifications, as well as bacteria classified as mitochondria or chloroplasts. Other problems included sequences classified as eukaryote taxa in the RDP and SILVA databases, and non-adherence to the binomial nomenclature in SILVA. As the research community continues to use these reference databases extensively, improved quality control will be necessary.

## Methods

### Collection of Prokaryote Nomenclature

Custom Python (v. 3.7.0) and bash shell scripts were created to collect and process reference nomenclature data from CyanoDB, LPSN, and Prokaryotic Nomenclature Up-To-Date. Nomenclature collected by the LPSN curators is available on the LPSN website (http://www.bacterio.net/). The Beautiful Soup (v. 4.6.0; https://www.crummy.com/software/BeautifulSoup/) Python module was used to scrape valid and invalid names from the LPSN website using CSS selectors (Additional file 1: Table S4) on July 5^th^, 2018.

Valid prokaryote names were obtained from the Prokaryotic Nomenclature Up-To-Date web service (https://bacdive.dsmz.de/api/pnu/) on July 5^th^, 2018. Multiple commonly used prokaryote names that are not validly published are listed in files available on the Up-to-Date website (https://www.dsmz.de/support/bacterial-nomenclature-up-to-date-downloads.html). The Up-to-Date xlsx file (last updated October 2017) was downloaded and converted to tab separated format using OpenOffice (v. 4.1.5). Invalid names were collected using names classified as ″orthographically incorrect name″, ″illegitimate name″, and ″rejected name″ in the status column.

Cyanobacteria nomenclature was obtained from the CyanoDB webpage (http://www.cyanodb.cz), which was last updated in April 2014. The webpages listing valid genera and their corresponding type species, as well invalid names were copied into text files, and parsed using Python scripts.

### Nomenclature Validation Set Curation

Pairwise comparisons between LPSN and Prokaryote Nomenclature Up-to-Date were performed to estimate the comprehensiveness and accuracy of the validation set (Additional file 1: Table S5, Table S6). The nomenclature in these databases was considered reliable, as the validity of 98% of the genera and 99% of species present in both databases was agreed upon. The final validation set contained three categories: valid names, invalid names, and disputed names. Names were considered valid if they were deemed valid in at least one reference database and were not deemed invalid by any other. Similarly, names were considered invalid if they were deemed invalid in at least one reference database and were not deemed valid by any other. Disagreements on the validity of names between databases were considered disputed.

### Validation of Public 16S rRNA Gene Reference Database Nomenclature

Three specialized 16S rRNA gene databases were evaluated in this study: Greengenes (v. 13_5), RDP (release 11, update 5), and SILVA (SSU Ref, v. 132). Delimiter separated flat files containing the taxonomic classifications for each reference sequence were downloaded from each database provider and custom Python and bash shell scripts were used to extract taxonomic classifications at the genus (Greengenes, RDP, SILVA) and species (Greengenes, SILVA) levels. Manual evaluations of the taxonomy files revealed metadata in the taxonomic classifications for all three databases. Therefore, lists of obvious metadata annotations were created for each database (Additional file 1: Table S1) and used to extract records annotated with metadata.

Bash shell scripts were created to categorize the 16S database nomenclature annotations using the validation set. The R Taxize package (v. 0.9.3) was used to query the unclassified annotations using the NCBI E-utilities API for taxonomic information. The Taxize results were used to identify taxa classieGreengenes, RDP, and SILVA all contained chloroplasts and mitochondria as taxonomic groups. Bash scripts were used to identify sequences classified as belonging to chloroplasts and mitochondria. Custom Python scripts were used to verify the correct usage of the binomial nomenclature for the species contained in Greengenes and SILVA.

## Declarations

### Ethics approval and consent to participate

Not applicable

### Consent for publication

Not applicable

### Availability of data and material

All scripts used in this study are available in the https://github.com/kyleLesack/16sdbnomenclature github repository. The results of this study are provided in the additional supporting files. The 16S rRNA gene reference database taxonomic classifications are available from the Greengenes (http://greengenes.secondgenome.com/), RDP (https://rdp.cme.msu.edu), and SILVA (https://www.arb-silva.de) websites. The prokaryote nomenclature reference data is available from the CyanoDB (http://www.cyanodb.cz/), LPSN (http://www.bacterio.net/), and Prokaryote Nomenclature Up-to-Date (https://www.dsmz.de/bacterial-diversity/prokaryotic-nomenclature-up-to-date) websites.

Additional file 1: **Table S1**. Keywords used to filter metadata from Greengenes and SILVA. **Table S2**. Most common invalid and uncategorized genus annotations in Greengenes, RDP, and SILVA. **Table S3**. Most common invalid and uncategorized species annotations in Greengenes, RDP, and SILVA. **Table S4**. CSS selectors used to scrape nomenclature data from LPSN. **Table S5**. Pairwise comparisons of genera in both databases. **Table S6**. Pairwise comparisons of species in both databases.

Additional files 2-3: Unique eukaryote names and records that were identified in the RDP taxonomic annotations.

Additional files 4-5: Unique eukaryote names and records that were identified in the SILVA taxonomic annotations.

Additional files 6-11: The accompanying csv files contain the valid, invalid, and unclassified genera and species contained in Greengenes. The files contain two columns: (1) the Greengenes sequence identification number, (2) the nomenclature annotations for the given rank.

Additional files 12-14: The accompanying csv files contain the valid, invalid, and unclassified genera contained in RDP. The files contain two columns: (1) the RDP sequence identification number, (2) the genus nomenclature annotations.

Additional files 15-20: The accompanying csv files contain the valid, invalid, and unclassified genera and species contained in SILVA. The files contain two columns: (1) the SILVA sequence identification number, (2) the nomenclature annotations for the given rank.

Additional files 21-25: The accompanying csv files contain taxa classified as either mitochondria or chloroplasts in Greengenes, RDP, and SILVA. The files contain the database identification numbers, and delimiter separated taxonomic classifications.

Additional files 26-28: The accompanying csv file contains the SILVA records that failed the binomial nomenclature test. The file contains three columns: (1) the SILVA identification number, (2) the genus name, (3) the species name.

### Authors’ contributions

The work presented here was designed and conducted by KL under the supervision of IB. KL wrote the scripts used to collect and analyze the data. The main text was written by KL with input IB.

## Acknowledgements

Not applicable

